# Unveiling Cas8 Dynamics and Regulation within a transposon-encoded Cascade-TniQ Complex

**DOI:** 10.1101/2024.06.21.600075

**Authors:** Amun C. Patel, Souvik Sinha, Pablo R. Arantes, Giulia Palermo

**Affiliations:** Department of Bioengineering, University of California Riverside, 900 University Avenue, Riverside, CA 52512, United States; Department of Chemistry, University of California Riverside, 900 University Avenue, Riverside, CA 52512, United States

## Abstract

The *Vibrio cholerae* Cascade-TniQ complex unveiled a new paradigm in biology, demonstrating that CRISPR-associated proteins can direct DNA transposition. Despite the tremendous potential of “knocking in” genes at desired sites, the mechanisms underlying DNA binding and transposition remain elusive. In this system, a conformational change of the Cas8 protein is essential for DNA binding, yet how it occurs is unclear. Here, structural modeling and free energy simulations reconstruct the Cas8 helical bundle and reveal an open-to-close conformational transition at key steps of the complex’s function. We show that when Cascade-TniQ binds RNA, the Cas8 bundle changes conformation mediated by the interaction with the Cas7.1 protein. This interaction alleviates unfavorable contacts and synchronizes Cas8’s shift with neighboring subunits, lowering the barrier for the transition to the open state, a critical requirement for DNA binding. As DNA fully pairs with RNA, the open state becomes increasingly accessible, favoring interactions with DNA and aiding the formation of an R-loop. These outcomes provide the first dynamic representation of a critical conformational change in one of the largest CRISPR systems and illustrate its role at critical steps of the Cascade-TniQ biophysical function, advancing our understanding of nucleic acid binding and transposition mechanisms.

---

DNA transposition is a fundamental process enabling the “jumping” of genes from one location to another in the genome. Recent evidence revealed a functional relationship between certain CRISPR (Clustered Regulated Interspaced Palindromic Repeats) systems and DNA transposition^1,2^, showing that bacteria have evolved RNA-guided programmable machinery to transpose genes. This has demonstrated a new paradigm in biology, where CRISPR proteins direct the activity of transposon enzymes to “knock-in” genes at the desired site^3^. The implications of this discovery for genome editing are profound and pave the way for targeted genomic insertion.

CRISPR-Cas systems use a guide RNA to recognize and cleave target DNA sequences^4^. Upon detection, these complexes conformationally and dynamically adjust to form an “R-loop” structure, whereby one DNA strand – the target strand (TS) – matches the guide RNA and the other non-target strand (NTS) is displaced^5^. The Cas proteins may then either directly cut the DNA or recruit an external nuclease. By design, such systems are ideal for “knocking out” genes, but the insertion of genes remains challenging. To site-specifically “knock-in” genes at the desired site, bacteria have evolved CRISPR-Cas variants lacking cleavage capability^1,2^. These variants originate from CRISPR-associated transposons (CASTs) and, upon identifying the target DNA, recruit key transposition proteins for site-specific DNA transposition. In *Vibrio Cholerae*, a Tn7-like transposon was found to encode a type 1F CRISPR-Cas system^1^ – also known as “Cascade” – a majestic multi-subunit complex that targets DNA through several Cas proteins (**Fig. 1a**)^6,7^. This system associates with a homodimer of the transposition protein TniQ to form a “Cascade-TniQ” complex. Cascade-TniQ then undergoes a search for the target DNA, followed by dynamic R-loop formation and the recruitment of the transposon for DNA integration through the aid of additional proteins, also encoded in the Tn7-like transposon. In this complex transposition machinery, R-loop formation is a key regulatory checkpoint. However, its mechanistic details and dynamic regulation remain highly ambiguous.

**Fig. 1.**
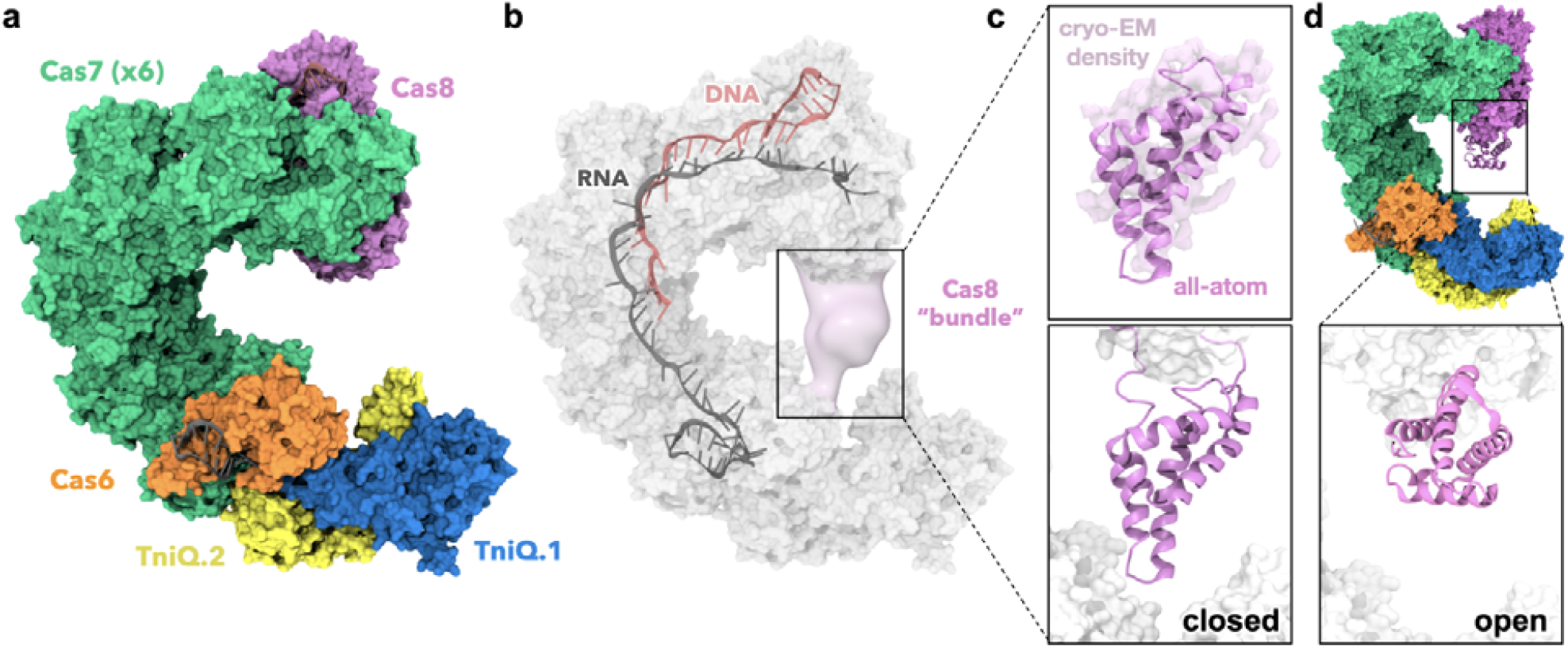
Overview of the *Vibrio Cholerae* Cascade-TniQ complex. **a**. Cryo-EM structure of the Cascade-TniQ complex bound to RNA and DNA (PDB: 6PIJ)^6^, consisting of several Cas proteins (Cas6, orange; six Cas7 subunits (green); Cas8, pink), and the TniQ dimer (TniQ.1, yellow; TniQ.2 blue). **b**. The structure captures an incomplete R-loop state, where the target DNA (salmon) partially binds the guide RNA (gray) with 20 base pair complementarities, rather than 32. The cryo-EM map (EMD-20351, 3.5 Å resolution) shows weak electron density for a crucial region of Cas8, termed the ‘Cas8 bundle’ (residues 277-385). **c-d**. Structural modeling of the Cas8 bundle, illustrating both the closed (**c**) and open (**d**) conformations. For the closed state (**c**, top panel), a comparison is shown between the all-atom model and the weak electron density for the Cas8 bundle in the EMD-20351 map.

Structures of the *V. Cholerae* Cascade-TniQ revealed that the complex comprises six Cas7 subunits that constitute the backbone of the structure, which accommodate the nucleic acids (**Fig. 1a**)^6,7^. One Cas7 protein (Cas7.1) and Cas6 locate at the head of the complex and bind the TniQ homodimer, while the Cas8 protein occupies the tail of the complex. An intriguing aspect is represented by the observation that a key structural element of the Cas8 subunit – a helical bundle composed of residues 278–388, referred to as the “Cas8 bundle” – could not be modeled from cryo-electron microscopy (cryo-EM) images of the *V. Cholerae* Cascade-TniQ complex during R-loop formation (**Supplementary Fig. 1**). Indeed, the electron density maps obtained by Halpin-Healy et al.^6^ show weak electron density for the Cas8 bundle in the same location before R-loop formation (i.e., in the RNA-bound state, EMD-20350 map) and in an incomplete R-loop state, in which the guide RNA-target DNA duplex is only partially formed (i.e., incomplete DNA-bound state, EMD-20351, **Fig. 1b**).

Structural studies on a related type 1F Cascade from *P. aeruginosa* have also indicated that the Cas8 bundle adopts distinct conformations at various stages of R-loop formation^8,9^. Moreover, recent cryo-EM data on 1F3-b, a structurally similar Cascade complex to the 1F3-a studied here, demonstrates a locked Cas8 bundle which binds the DNA upon complete heteroduplex formation^10^. This occurs as the DNA TS rejoins the NTS, reinforcing the notion that a conformational change in the bundle is essential for nucleic acid binding. However, this structural transition and its implications before and after R-loop formation remain elusive.

Here, extensive structural modeling and free energy simulations of the Cascade-TniQ complex are used to establish the structure and dynamics of the Cas8 helical bundle at different stages of R-loop formation. We characterize two main configurations of the Cas8 bundle, open and closed, and detail the conformational transition between them. We show that in the RNA bound states, the Cas8 bundle undergoes a conformational shift facilitated by its interaction with the Cas7.1 protein. An in-depth analysis of the complex dynamics through wavelet decomposition shows that this interaction aligns Cas8’s movements with adjacent subunits, easing the transition to the open state. As DNA pairs more fully with RNA, the open state becomes increasingly achievable, promoting interactions with DNA and supporting R-loop formation. Hence, we identify two critical contributors to the Cas8 conformational change during the R-loop formation process, a key Cas8-Cas7.1 interaction, and the RNA-DNA duplexing. These findings offer the first dynamic depiction of an essential conformational shift within one of the largest CRISPR systems, deepening our understanding of its biophysical process and transposition mechanisms.

## Results

### Cas8 binding mode

To describe the conformation of the Cas8 bundle bound to the Cascade-TniQ complexes, we performed structural modeling, followed by molecular dynamics (MD)-driven cryo-EM fitting. Alpha-fold2^11^, an Artificial Intelligence (AI)-powered method that incorporates biophysical knowledge about protein structure and multi-sequence alignments, was used to predict the three-dimensional structure of Cas8. AlphaFold2 was combined with PyRosetta^12^, preserving the positions of the experimental structures, while adding missing loops and the Cas8 bundle from the AlphaFold2 model (details in the Methods section).

Structural modeling reported two main configurations of the Cas8 bundle – i.e., closed and open states (**Fig. 1c-d, Supplementary Figs. 2-3**) – that resemble the available experimental data. In detail, the open state of the Cas8 bundle (**Fig. 1d)** resembles the orientation observed in the type 1F Cascade from *P. aeruginosa* obtained by Guo and co-workers (PDB: 6B45)^8^. In the closed state (**Fig. 1c**), the Cas8 bundle assumes a conformation similar to the weak electronic density observed by Halpin-Healy and coworkers (EMD-20350, EMD-20351 maps^6^). Furthermore, the closed state is evidenced in the top ranked AlphaFold2 model, followed by the open state, displaying a comparable level of confidence in the prediction (**Supplementary Fig. 3**, details in the Methods section). This indicates that both states are equally possible and further reinforces the concept of a conformational change between them ^8,9^.

To refine the position of the closed Cas8 bundle with respect to the weak cryo-EM density observed by Halpin-Healy, we used a molecular dynamics flexible fitting (MDFF) approach^13,14^. The cryo-EM maps display low-resolution density (from ∼6 to 9 Å) at the level of the Cas8 bundle^6^. Using these cryo-EM maps as a target, we carried out a flexible fitting of our computational model into the experimental density. The fitted structures refined the missing loops and the Cas8 bundle with respect to the experimental densities (**Supplementary Fig 4**). In the final structures, the Cas8 bundle binds with its apical portion the Cas7.1 and TniQ.2 proteins, reflecting the experimental data (**Fig. 1c**). It is also notable that the fitted structure of the Cas8 bundle aligns with a poly-alanine model reported by Li and colleagues, derived from weak electron density data (**Supplementary Fig 5**)^15^.

### Conformational change of the Cas8 bundle

To describe the conformational change between the closed and open states of the Cas8 bundle, we performed free energy simulations using the Umbrella Sampling (US) method^16^ (details in Supplementary Methods). The structural transition was analyzed in the available cryo-EM structures captured prior to R-loop formation (RNA-bound) and in the DNA-bound state, obtained by Halpin-Healy (HH)^6^ and Jia^7^. Specifically, the RNA-bound systems were based on PDB 6PIG (referred to as RNA-HH) and 6V9Q (RNA-Jia). For the DNA-bound states, we considered PDB 6PIJ (DNA-HH) in which the R-loop is partially formed (i.e., the target DNA binds the guide RNA with 20 base pair complementarities, rather than 32), and 6VBW (DNA-Jia) that displays a formed R-loop (**Supplementary Fig. 1**). The open-to-closed transition (and *vice versa*) was rigorously investigated by sampling the populations of the two states and through converged US simulations (**Supplementary Fig. 6**). The transition was sampled along the difference in root-mean-square deviation of the heavy atoms’ positions in the Cas8 bundle (Cas8 RMSD) relative to both states, used as the reaction coordinate (RC).

The free energy profiles for the RNA-bound complexes reveal that in the system structured by Halpin-Healy (RNA-HH), the Cas8 bundle is trapped in a closed state minimum and lacks a stable open state (**Fig. 2a**). In the Cascade-TniQ structured by Jia (RNA-Jia), the Cas8 bundle displays a minimum corresponding to the closed state, and a diffusive region leading to a fully open state.

**Fig. 2.**
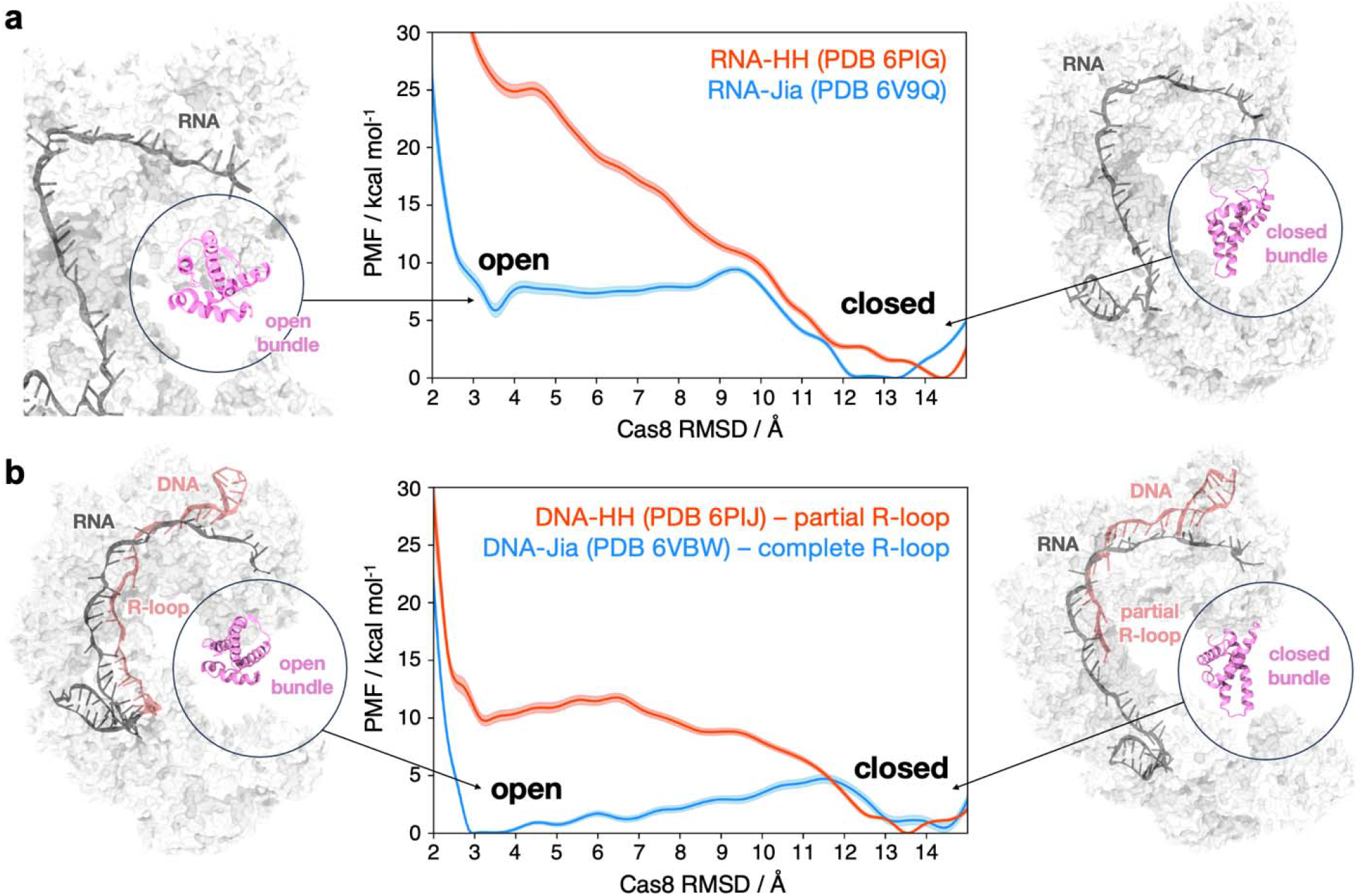
Conformational change of the Cas8 bundle from an open to a closed state. Free energy profiles for the Cas8 bundle conformational transition in the RNA-bound (**a**) and DNA-bound (**b**) Cascade-TniQ complexes computed through Umbrella Sampling simulations^16^. The difference in root-mean-square deviation of the heavy atoms’ positions in the Cas8 bundle, relative to both open and closed states (Cas8 RMSD), was used as the reaction coordinate. The free energy profiles were derived through the Variational Free Energy Perturbation (vFEP) method^17^, using bootstrap analysis to compute the associate errors (details in Supplementary Methods). **a**. The RNA-bound systems were based on PDB 6PIG (referred to as RNA-HH) and 6V9Q (RNA-Jia). In the RNA-HH system, the Cas8 bundle is trapped in a closed state (shown on the right). In RNA-Jia, the Cas8 bundle shows diffusive region leading to a fully open state (left). **b**. The DNA-bound systems were based on PDB 6PIJ where the R-loop is partially formed (DNA-HH, right), and PDB 6VBW, displaying a fully formed R-loop (DNA-Jia, left). Upon partial R-loop formation (DNA-HH), the Cas8 bundle prefers the closed state (shown on the right), whereas upon complete formation of the R-loop (DNA-Jia), the open and closed states are separated by ∼5 kcal/mol.

In the DNA-bound state, the complex obtained by Halpin-Healy (DNA-HH) displays a higher barrier between closed and open states (i.e., ∼10 kcal/mol), compared to Jia’s complex (DNA-Jia), in which a ∼5 kcal/mol barrier separates the two states (**Fig. 2b**). Notably, the Halpin-Healy structure is representative of an incomplete R-loop state, whereas the R-loop is formed in Jia’s structure. Hence, when the R-loop is partially formed, the closed state is favored, whereas upon R-loop formation, the bundle assumes both the open and closed states, separated by a ∼5 kcal/mol barrier. This indicates that, as the DNA TS binds the RNA forming the R-loop, it lowers the barrier for accessing the open state. This is in line with the hypothesis, based on the biochemical data of a related type 1F Cascade from *P. aeruginosa*, that conformational changes of the bundle could aid R-loop formation^8,9^. In their paper, Rollins and colleagues demonstrated that the substitution of the positively charged residues in the bundle impairs DNA binding^9^, suggesting that conformational changes in this region may facilitate R-loop formation by interacting with the DNA. Our findings further support this hypothesis, showing that as the R-loop forms, the open state of the bundle becomes more thermodynamically accessible. As the Cas8 bundle transitions to an open state, it exposes positively charged residues, essential for interacting with DNA. Hence, the ease of the Cas8 conformational change can facilitate the interaction between the bundle and DNA, ultimately promoting R-loop formation.

### Cas7.1 favors the conformational change of the Cas8 bundle in the RNA-bound complex

Free energy simulations revealed an appreciable discrepancy in the free energy profiles of the RNA-bound Cascade-TniQ complexes (**Fig. 2a**). To understand this observation, we analyzed the interactions established by the Cas8 bundle along its conformational change. As a result, the bundle interacts predominantly with the Cas7 subunits (**Supplementary Fig. 7)**. In Jia’s RNA-bound system, it mainly interacts with Cas7.1, while it prefers Cas7.2 in the HH’s system. Interestingly, a loop of approximately 10 amino acids within the Cas7.1 subunit (residues 51-60, hereafter referred to as Cas7.1 loop) exhibits a notable difference in position relative to the Cas8 bundle. In Jia’s RNA-bound system, when the Cas8 bundle assumes a closed state, this Cas7.1 loop binds the Cas8 bundle (**Fig. 3a, Supplementary Fig. 7**). In contrast, this interaction is absent in HH’s RNA-bound complex (**Fig. 3b**).

**Fig. 3.**
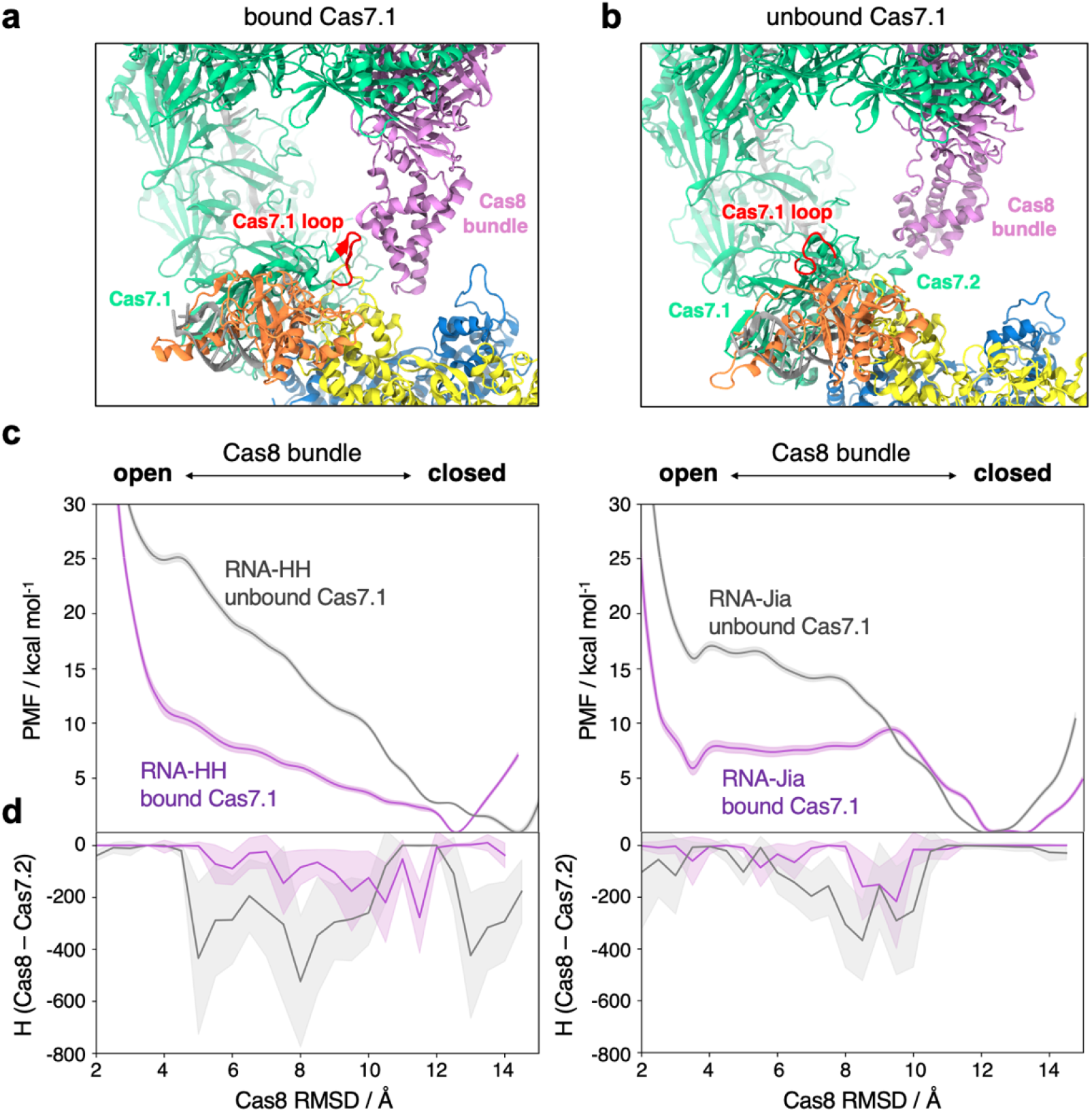
Role of Cas7.1 in the Cas8 conformational change. **a-b**. Bound (**a**) and unbound (**b**) conformations of the Cas7.1 domain, showing the Cas7.1 loop (residues 51-60, red) either bound to Cas8 or distant from it. **c**. Free energy profiles for the Cas8 bundle open-to-close conformational change in the RNA-bound systems (RNA-HH, left; RNA-Jia, right), in the presence of a bound (purple) and unbound (gray) Cas7.1. The free energy profiles were computed along the difference in RMSD of the Cas8 bundle heavy atoms, relative to both states (Cas8 RMSD). Bootstrap analysis was applied to compute the associate errors (details in Supplementary Methods). **d**. Proxy of enthalpy between Cas8 and Cas7.2, (Cas8 – Cas7.2), computed as the product of the number of contacts and the interaction energies along the reaction coordinate. Errors are computed as the standard deviation per window (details in Supplementary Methods).

We thereby sought to understand the importance of Cas7.1 binding on the conformational change of the Cas8 bundle. Toward this aim, we performed additional free energy simulations of the RNA-HH and the RNA-Jia systems in the presence of a Cas7.1 bound and unbound to Cas8 in the closed state. The US approach was applied to simulate the open-to-closed conformational transition of Cas8, as described above (details in Supplementary Methods, **Supplementary Fig. 8**). These simulations report a reduction in the free energy profile in the RNA-HH system when Cas7.1 binds Cas8, compared to the system with an unbound Cas7.1 (**Fig. 3c**). A similar trend is evident in the RNA-Jia system, where the free energy profile decreases in the presence of a bound Cas7.1 (**Fig. 3d**). This indicates that, in the RNA-bound state, the Cas7.1-Cas8 interaction favors the conformational change of Cas8 bundle.

Analysis of the interactions between the Cas8 bundle and the Cas7 subunits along the free energy profiles reveals that when Cas7.1 is unbound, the interactions between Cas8 and Cas7.2 are stronger than when Cas7.1 is bound (**Supplementary Figs. 7, 9**). Indeed, when the Cas7.1 loop does not interact with Cas8, the bundle binds the hairpin loops of Cas7.2 (residues 48-63, **Fig. 3b**), hampering its transition. This explains why the Cas8 bundle is trapped in a minimum corresponding to a stable closed conformation. To investigate whether interactions between the bundle and hairpin loops of Cas7.2 contribute to the observed discrepancies in free energy profiles, we computed a proxy for enthalpy. Specifically, we measured the product of the number of contacts and interaction energies between the two regions of interest (residues 278-388 of Cas8 and residues 48-63 of Cas7.2), normalized by the average number of contacts throughout the simulations (details in Supplementary Methods).

Values approaching zero indicate minimal interaction, while larger absolute values reflect stronger or more extensive interactions. In the presence of an unbound Cas7.1, our enthalpic proxy increases (in absolute terms) as the Cas8 bundle changes conformation (**Fig. 3d**). This shift corresponds to stabilizing interactions with the hairpin loops of Cas7.2 in both the HH and Jia RNA-bound systems. Notably, these enhanced interactions are associated with steeper regions in the free energy profiles (**Fig. 3c**). This indicates that, in the presence of an unbound Cas7.1, the interactions between the Cas8 bundle and Cas7.2 play a critical role in shaping the free energy profiles and disfavor the transition to an open state. In contrast, when Cas7 is bound, these interactions are minimal and do not significantly influence the free energy profiles. Overall, these additional free energy simulations indicate that, in the RNA-bound states, the interaction with Cas7.1 promotes the Cas8 transition, mitigating unfavorable contacts with Cas7.2.

### Synchronized dynamics reinforce the role of Cas7.1

Cas7.1 plays a critical role in the open-to-closed conformational transition of the Cas8 bundle within the RNA-bound Cascade-TniQ complex (**Fig. 3**). To understand how Cas8 dynamically couples with the other neighboring protein subunits (i.e., Cas6, and TniQ) during the open-to-close conformational change, we performed a wavelet analysis^18^. This analysis decomposes the trajectories of the atoms of interest (i.e., the C⍰ of Cas8, Cas6, and TniQ) into wavelets to examine the coupling between the wavelet scales of different domains. Wavelet heat maps have been built by plotting the wavelet scale for the Cascade-TniQ domains of interest along the open-to-close transition of the bundle (**Fig. 4**). Short wavelet scales capture fast protein motions, while larger scales correspond to slower motions (details in the Methods section).

**Fig. 4.**
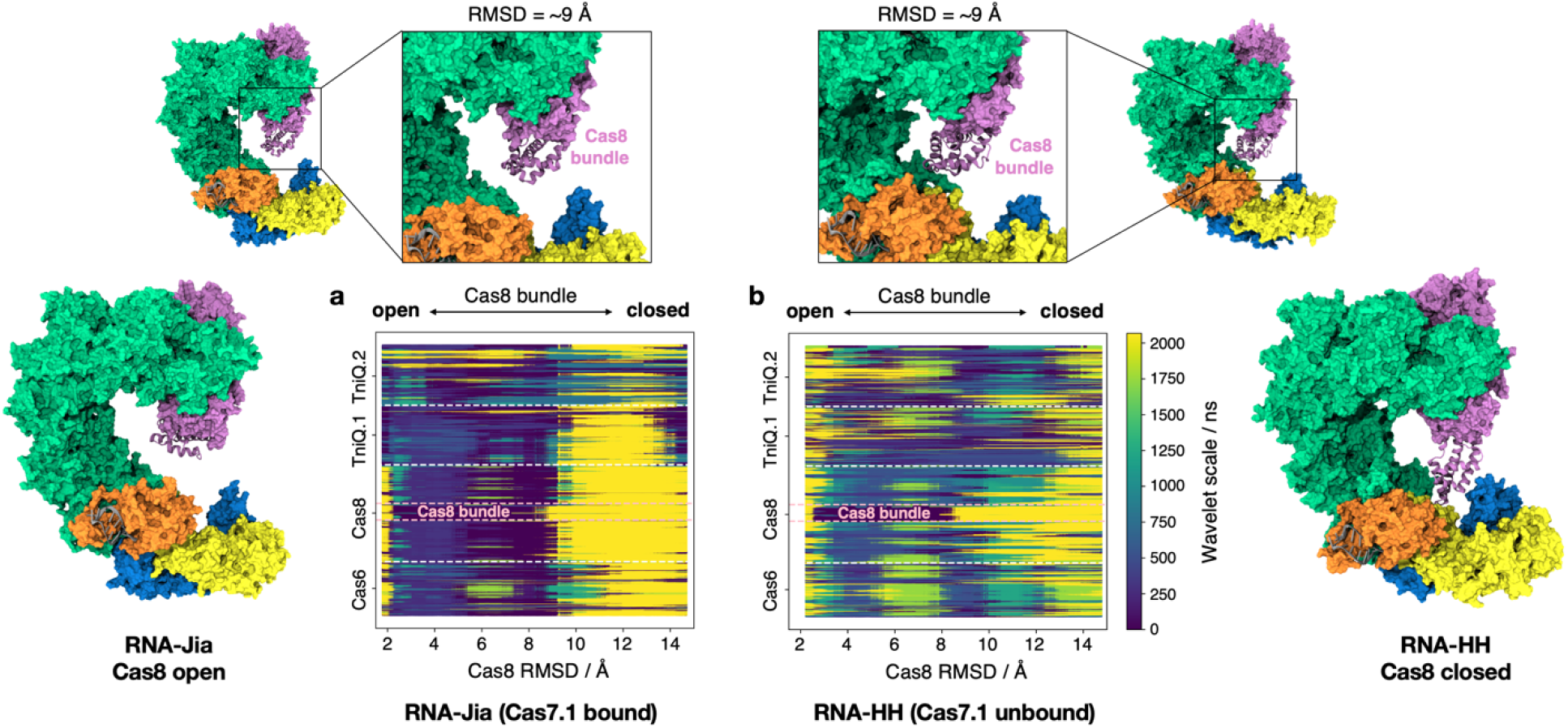
Coupling of Cas8 conformational changes with the dynamics of neighboring protein subunits. Wavelet maps decompose the system’s dynamics across various time scales, distinguishing between long-scale (low-frequency, yellow) and short-scale (high-frequency, blue) vibrational wavelets. Wavelet analysis was conducted on Cas8 and its neighboring subunits (i.e., the C⍰ of Cas8, Cas6 and TniQ.1-2) along the open-to-close conformational transition of the Cas8 bundle in the RNA-bound systems, where the Cas7.1 loop is either bound (RNA-Jia, **a**) or unbound (RNA-HH, **b**) to Cas8. This analysis was performed along the reaction coordinate (Cas8 RMSD) of umbrella sampling simulation (details in the Methods section). In RNA-Jia, when Cas7.1 is bound to Cas8, we observe a distinct shift from fast to slow modes as the Cas8 bundle assumes a closed state (at ∼9 Å RMSD), while in RNA-HH, a mix of modes in the wavelet scales is observed. Cartoons of the Cascade-TniQ complex are depicted at various values of the reaction coordinate (Cas8 RMSD) to illustrate the conformations adopted by the system.

In Jia’s RNA-bound complex, in which Cas7.1 is bound to Cas8, we observe a distinctive shift from fast to slow modes as the bundle falls into the closed state (at ∼9 Å RMSD, **Fig. 4a**). This shift couples the dynamics of Cas8 with Cas6, and TniQ. In the RNA-HH system, where Cas7.1 is unbound, the system acquires a cacophonous mix of modes in the wavelet scales (Fig. 4b) and as the bundle changes conformation, its dynamics decouple from those of the neighboring domains. This trend is maintained in the RNA-HH and RNA-Jia systems displaying a bound and unbound Cas7.1, respectively (**Supplementary Fig. 10**) and demonstrates an important role of TniQ and Cas6 during the Cas8 conformational change. Since the Jia and HH’ systems mainly differ in the Cas7.1 gate orientation, these analyses reinforce the role of Cas7.1 during the bundle conformational change, synchronizing the motions of Cas8 with its neighboring Cas7.6 and TniQ.

Collectively, our findings indicate that Cas7.1 facilitates the open-to-closed conformational shift in the Cas9 bundle by serving two distinct functions. First, it reduces unfavorable interactions with Cas7.2 (**Fig. 3**), contributing an enthalpic effect. Second, it dynamically couples the Cas8 subunit with Cas7.6 and TniQ (**Fig. 4**), driving a coordinated shift in the conformational states of these subunits.

## Discussion

The *Vibrio cholerae* Cascade-TniQ complex has introduced a new paradigm in biology, demonstrating that CRISPR-associated proteins can guide the precise transposition of genes to targeted sites^1,2^. Structural studies of the Cascade-TniQ complex, captured at various stages of R-loop formation, did not resolve the highly dynamic Cas8 bundle^6,7^. This motif was shown to play a role during R-loop formation in similar systems^8,9^, raising questions about how its conformational changes influence the binding of target DNA to the guide RNA. Thus, we aimed to elucidate the dynamics of the Cas8 bundle within the Cascade-TniQ complex. Structural modeling identified two main configurations of the bundle – open and closed states – that align with existing structural studies and suggest an open-to-closed conformational transition, which we thoroughly investigated using free energy methods.

Studies of the conformational transition in the RNA-bound states reveal that the Cas8’s shift is facilitated by its interaction with Cas7.1, binding a 10 residues loop (residues 51-60). This interaction mitigates unfavorable contacts with Cas7.2 that would otherwise lock the bundle in a closed state (**Figs. 2-3**). Moreover, when Cas8 binds to Cas7.1, the conformational change of the bundle couples with the dynamics of the adjacent Cas7.6 subunit and the TniQ dimer (**Fig. 4**). This reinforces the role of Cas7.1, synchronizing the motions of Cas8 with its neighboring domains. Cas7.1 thereby exerts a dual critical role: it alleviates unfavorable contacts, yielding an enthalpic benefit, while also driving an entropic effect that synchronizes Cas8’s shift with neighboring subunits. This coordination effectively lowers the barrier for transitioning to the open state, which primes the complex for DNA binding.

In DNA-bound states, the Cas8 bundle undergoes a favorable conformational change as the DNA fully complements the guide RNA, forming the R-loop (**Fig. 2b**). Indeed, when the R-loop is only partially formed, the bundle preferentially adopts a closed conformation. As the R-loop forms, the open state becomes more accessible, facilitating the interaction between the positively charged residues of the bundle and the DNA. In this respect, biochemical experiments of a related type 1F Cascade from *P. aeruginosa* have demonstrated these interactions to be essential for DNA binding^9^. This suggests that, as the R-loop proceeds, a conformational change of the Cas8 bundle is critical to enable DNA binding and drive the process to completion.

In summary, integrating our findings with existing experimental evidence enables us to propose a mechanistic model outlining the role of the Cas8 bundle in nucleic acid binding (**Fig. 5**). In the RNA-bound state, the interaction between the Cas8 bundle and the Cas7.1 loop promotes the transition to the open state (**Fig. 5, path 1**). This opening is essential for DNA binding. Upon DNA binding, the open state becomes more accessible as the DNA target strand matches the guide RNA (**Fig. 5b**). The opening of the helical bundle at this stage is essential to enable its positively charged residues to interact with the DNA, facilitating its repositioning as suggested by Rollins and colleagues^9^. This leads to a completely formed R-loop (**Fig. 5c**). On the other hand, in the absence of a Cas8-Cas7.1 interaction (**Fig. 5, path 2**), the Cas8 bundle cannot easily transition to the open state (**Fig. 5d**), which inhibits full R-loop formation.

**Fig. 5.**
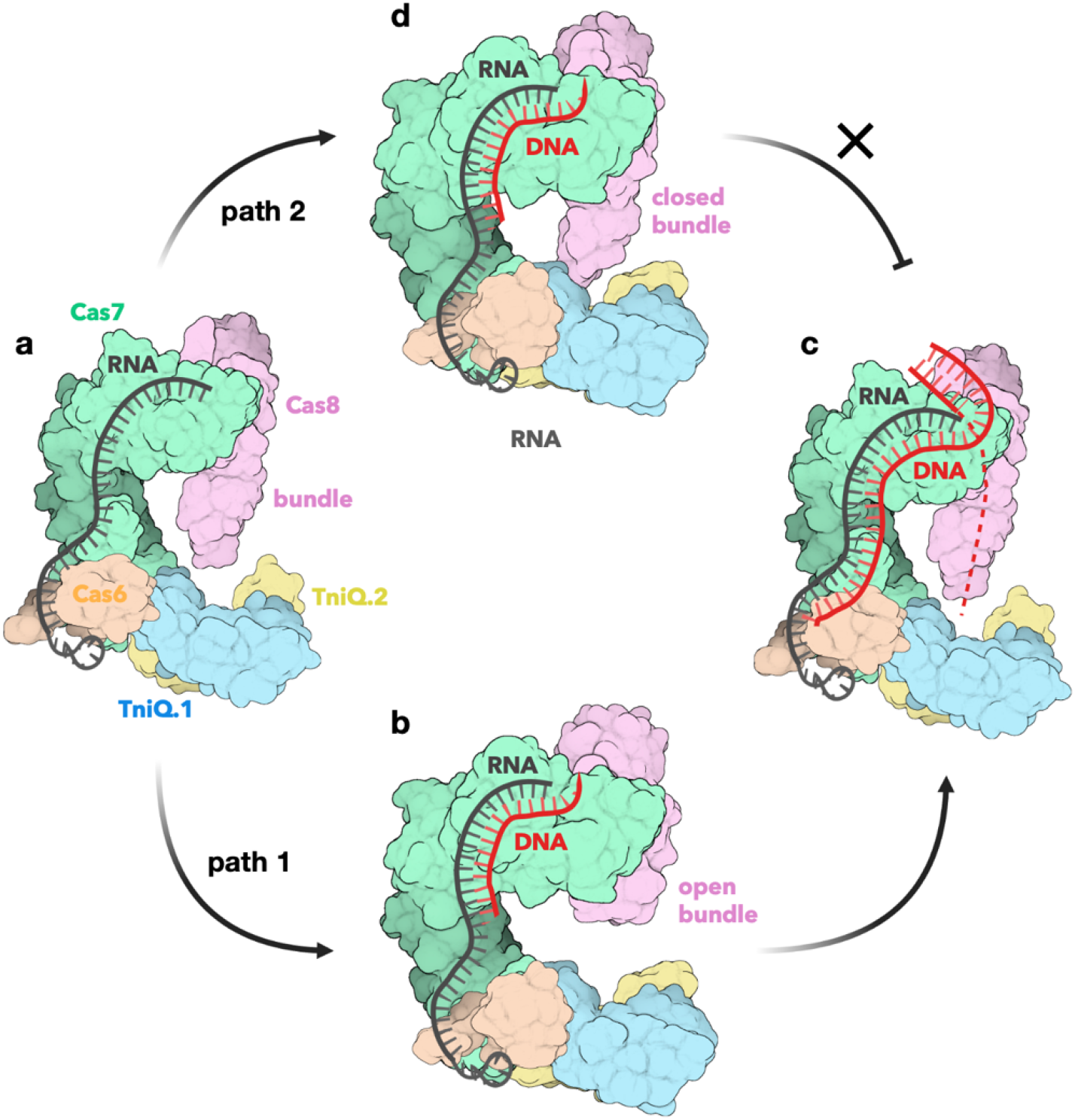
Model for the role of Cas8 in R-loop formation. In the RNA-bound state (**a**), the interaction between the Cas8 helical bundle and the Cas7.1 loop drives the transition of the bundle to the open state (path 1). Once the DNA target strand aligns with the guide RNA, the open state becomes more accessible (**b**). At this stage, the opened helical bundle facilitates the fully formed R-loop (**c**). Conversely, without the Cas8-Cas7.1 interaction (path 2), the Cas8 bundle does not easily transition to an open state (**d**), preventing complete R-loop formation.

The Cas8 conformational change is thereby critical to lead R-loop formation to completion. In this respect, it is insightful to compare our findings with other CRISPR-Cas systems. In the Cas proteins, the formation of the R-loop is facilitated by highly flexible recognition domains^19–21^. Notably, CRISPR-Cas systems do not utilize ATP hydrolysis for R-loop formation; instead, the dynamics of the conformational changes in their flexible subunits promote R-loop formation. This leads to a general theme at play in CRISPR-Cas systems, where DNA binding is coupled with large conformational changes of the binding domains.

Overall, these findings depict a highly dynamic helical bundle in the Cas8 protein, playing an essential role in R-loop formation. These outcomes provide the first dynamic and thermodynamic representation of a critical conformational change within one of the largest CRISPR systems and lay the foundation for further experimental investigations to fully elucidate Cas8’s role in R-loop formation. Overall, the insights into Cas8 dynamics presented here enhance our understanding of the transposon-encoded Cascade-TniQ biophysical mechanism and provide valuable contributions to our knowledge of the transposition process.

## Methods

### Molecular Simulations

Computational studies have been based on the cryo-EM structures of the *V. Cholerae* type 1F Cascade variant (referred to as the Cascade-TniQ system) in complex with RNA and DNA, as reported in two independent studies. Halpin-Healy et al.^6^ reported RNA-(EMD-20350, PDB 6PIG) and DNA-bound (EMD-20351, PDB 6PIJ) complexes at resolutions of 3.50 Å and 2.90 Å, respectively. Jia et al.^7^ obtained structures for the RNA-(EMD-21126, PDB: 6V9Q) and DNA-bound (EMD-21146, PDB: 6VBW) complexes at 2.90 Å and 3.20 Å, respectively. After structural modeling of the missing residues and the Cas8 bundle (vide infra), the structures were embedded in explicit water with Na^+^ ions added to neutralize the total charge under physiological conditions. Orthorhombic periodic simulation cells measuring 187 × 216 × 248 Å^3^ were used, resulting in approximately 928,000 atoms for each system. These systems have undergone molecular dynamics (MD) simulations using a protocol tailored for protein/nucleic acid complexes^22^, also applied in studies of CRISPR-Cas systems^23–25^. The Amber ff19SB^26^ force field was employed, incorporating the OL15^27^ and OL3^28^ corrections for DNA and RNA, respectively. The TIP3P^29^ model was used for water molecules. An integration time step of 2 fs was applied. The GPU-empowered version of AMBER 22^30^ was used as a MD engine. Full details on MD and free energy simulations are reported in the Supplementary Methods.

### Structural Reconstruction

The cryo-EM maps of the *V. Cholerae* Cascade-TniQ complex (EMD-20350, EMD-20351)^6^ obtained by Halpin-Healy et al. display weak electronic density (from ∼6 to 9 Å resolution) at the level of the Cas8 bundle (residues 277-385). To model the Cas8 bundle and reconstruct the missing loops (i.e., ∼10 amino acids within the Cas proteins), we employed AlphaFold2^11^ through the ColabFold^31^ notebook environment.

AlphaFold2 was applied using as input the amino acid sequence and an HMMer^32^ multiple sequence alignment method (default in the AlphaFold2 pipeline). This step aims to learn a rich pairwise representation that is informative about which residue pairs are close in 3D space. Then, this pairwise representation is used to directly produce atomic coordinates by treating each residue as a separate object, predicting the rotation and translation necessary to place each residue, and ultimately assembling a structured chain. Since AlphaFold2 is well-validated up to ∼1400 amino acid residues, we used this method to construct three partial multimers (MM) of the Cascade-TniQ system: MM1 composed of the Cas7.2–Cas7.4 proteins, MM2 composed of Cas8, Cas7.5 and Cas7.6, and MM3 composed of Cas7.1, Cas6, TniQ1 and TniQ2 (**Supplementary Fig. 2**). The accuracy of the obtained models was evaluated using a per-residue measure of local confidence (i.e., a predicted Local Distance Difference Test, pLDDT) on a scale from 0 (low confidence) to 100 (high confidence)^11^. A second metric, the Predicted Aligned Error (PAE), reports AlphaFold2’s expected position error at residue *x*, when the predicted and actual structures are aligned on residue *y*. Low values of PAE suggested that the AlphaFold’s structures exhibit confidence about the relative domain positions. The top-ranked model of the MM2 multimer displays a closed conformation of the Cas8 bundle (**Supplementary Fig. 2a**), similar to the weak electronic density observed by Halpin-Healy and co-workers in the RNA-bound state (EMD-20350 map, **Fig. 1c**)^6^. The MM2 multimer shows high pLDDT values for the ⍰-helical regions (**Supplementary Fig. 2c**), which also display low PAE values as a sign of high confidence in their relative position. Increased values of PAE are found for flexible loops within the Cas8 bundle, in line with the notion that the Cas8 bundle can adopt multiple conformations^8,9^. We then analysed the five top-ranked models of MM2 obtained by AlphaFold2, focusing on Cas8 and its neighbouring Cas7.6 (**Supplementary Fig. 3**). As a result, the first top-ranked model displays a close conformation of the Cas8 bundle, and the subsequent models also display an open conformation, similar to the orientation observed in the type 1F Cascade from *P. aeruginosa* obtained by Guo and co-workers (PDB: 6B45)^8^. Notably, both conformations have similar pLDDT and PAE values, indicating comparable levels of confidence in the predictions for both models.

The obtained AlphaFold2 models were used in conjunction with the cryo-EM structures to produce more accurate structural models of the Cascade-TniQ complexes. In detail, given the experimental structures and the AlphaFold2 models, the latter positions were threaded onto the former structures using PyRosetta^12^ with the RosettaCM^33^ threading mover. Importantly, we verified that the final RMSD between the cryo-EM structures and the corresponding threaded structures was zero. This preserved the position of the experimental structures, while also adding the missing loops and the Cas8 bundle from the AlphaFold2 model.

### Flexible Fitting into Low-Resolution Cryo-EM Density

To refine the position of the Cas8 bundle in the closed state with respect to the weak cryo-EM density observed by Halpin-Healy^6^, we used a molecular dynamics flexible fitting (MDFF) approach^13,14^. The cryo-EM maps obtained by Halpin-Healy display low-resolution density at the level of the Cas8 bundle (from ∼6 to 9 Å resolution)^6^. Using these cryo-EM maps as a target, we performed MD flexible fitting of our computational models, displaying a closed conformation of the Cas8 bundle. Specifically, our models of the RNA-bound and DNA-bound states were fitted into the respective cryo-EM densities (EMD-20350 and EMD-20351^6^) using MDFF. This approach applies an external biasing potential, *U*_*EM*_, where forces generated from the gradient of the EM density guide atoms toward areas of higher density.

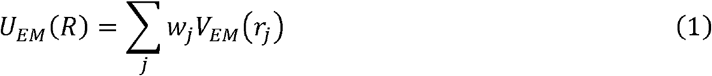

where *R* collects all atom coordinates and

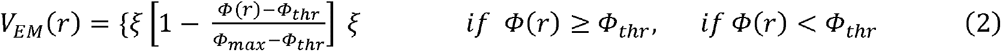

here, Φ(*r*) is the Coulomb potential from the EM density at position *r*, Φ_*max*_ is the maximum value of the EM density map, ξ is a force scaling factor, Φ_*thr*_ is a density threshold used to disregard the solvent density, *w*_*j*_ is a weight varied according to the type of atom placed in this potential (the atomic mass) and *r*_*j*_ is the position of atom *j*. The force 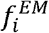 applied to an atom inside the potential is defined as:

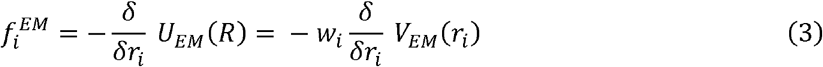

and can be tuned via a constant scaling factor ξ (same for all atoms), and the weight *w*_*j*_ that is defined on a per-atom basis. To preserve the integrity of secondary structures, restraints on bonds, angles, and dihedral angles were applied to regions with well-defined secondary structures, as identified by the Dictionary of Protein Secondary Structure (DSSP)^34^. In this work, the MDFF bias was applied in stages with progressively higher values of the scaling factor ξ in the range 0.01 to 0.1. Convergence was evaluated by assessing the root-mean-square deviation (RMSD) relative to the initial structure and the cross-correlation coefficient between the original EM density and a computed density map derived from the atomic model, following the approach described by Trabuco et al.^13^ Full details are reported in the Supplementary Methods.

#### Wavelet Analysis

Wavelet analysis decomposes a signal into wavelets, simplifying its structure similarly to Fourier series. The wavelet transform, widely used in data mining and applied by Heidari et al. to MD simulations^18^, breaks down a signal using orthogonal basis functions and generates wavelet coefficients. In this analysis, Continuous Wavelet Transforms (CWT) are used to decompose a time series into different coefficients characterized in the time scale domain. By further conducting a significance test, motions are classified as random or significant, identifying motions characterized by long-scale (low frequency) and short-scale (high frequency) vibrational wavelets. Wavelet analysis is performed on MD trajectories considering the displacement of each atom from a reference structure as an input signal. Here, the input signal was defined as the distance, in each frame, of atom from its position with respect to the first frame. To remove the global translational and rotational motions of the molecule, all frames were RMS-fit to the initial frame.

Wavelets, *w*(*t*), are defined by two parameters: the scale *S*_*k*_, indexed by the integer *k*, which stretches or compresses the wavelet, and translation, *j*, which shifts it along the time axis, resulting in an orthonormal family of wavelets, {*w*(*S*_*k*_ *t* − *j*)}_*j,k*∈*z*_. This family can be used to decompose any square integrable function. CWT are then used to reconstruct the original signal by continuously varying the parameters, capturing both frequency and temporal information. In detail, CWT allows for the application of wavelets continuously over time via convolution of the signal *r*(*t*) with the wavelet *w*_*k*_(*t*), where the translation parameter, *j*, is incorporated through convolution (denoted by the ^*^ symbol). For each atom, CWT is calculated by convolving the displacement data (i.e., the input signal) with the selected wavelet using a Fourier transform (FT) algorithm. Here, Morlet wavelets are applied, as by Heidari et al ^18^. CWT for atom *i* at scale *k* are calculated as follows:

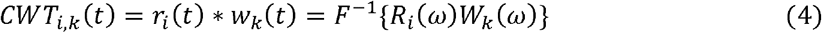

where *r*_*i*_(*t*) and *R*_*i*_(ω) denote the displacement trajectory for atom *i* in the time and frequency domains, respectively. *w*_*k*_(ω) and *w*_*k*_(ω) are the wavelet function at scale *k* in the time and frequency domains, respectively. *F* is the Fourier transform (inverse shown as *F*^−1^), and the ^*^ sign refers to convolution. Through this equation, CWT are computed using Morlet wavelets,, for each atom, repeated for all scale values indexed by k (i.e., *k* = 0, 1, … …). The scale *S*_*k*_ values are defined as *S*_*k*_ = *S*_0_ 2^*k*·*ds*^, where d, represents the interval between discrete scales. To satisfy the Nyquist-Shannon sampling theorem^35^, *ds*, is constrained by the sampling frequency to be at least twice the separation (in ns) between adjacent frames in the simulation, preventing aliasing. Here, trajectories were saved at 0.5 ns intervals; thus, *ds*, was set to 1. Consequently, *S*_0_ = 2 and 11 scales were used, indexed from *k* = 0 to *k* = 10, which generated a distribution of values *CWT*_*i,k*_(*t*) across scales *S*_*k*_.

Since this distribution is hypothesized to be a normal distribution, significance testing can be applied to determine the scale at which the greatest deviation from normality occurs. This enables us to pinpoint the scales where the atom exhibits significant motion. To simplify the significance testing of wavelet functions, the absolute values of the wavelet coefficients were used. Specifically, the distribution of the squared absolute values of the wavelet coefficients, ❘*W*_*i,k*_❘^2^, follows a chi-square distribution, as described below:

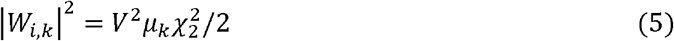

here, *V*^2^ is the *i*^*th*^ atom’s distance variance, 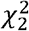 is the chi-square distribution with two degrees of freedom, is the mean of the expected distribution, given by μ_*k*_ = 0.002527 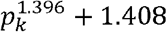, and p_*k*_ = 1.01. S_*k*_ for Morlet wavelets ^18^. The expected distribution of the absolute square of the wavelet coefficients, ❘*W*_*i,k*_❘^2^, is compared to the actual square of the wavelet coefficients ❘*CWT*_*i,k*_❘^2^.

If the coefficients fail the statistical test, then, ❘*CWT*_*i,k*_❘^2^ < ❘*W*_*i,k*_❘^2^,and these scales are the result of random noise and are excluded from further analysis. If all coefficients fail at a particular time point, the output defaults to zero. For the coefficients that pass the test, the scale with the greatest deviation from the expected chi-square distribution, specific to each atom and time point, is visualized using a heat map. Here, wavelets analysis was performed on Cas8 and its neighboring subunits (i.e., the C⍰ of Cas8, Cas6 and TniQ.1-2) along the open-to-close conformational transition of the Cas8 bundle in the RNA-bound systems. This analysis was conducted along the reaction coordinate of the umbrella sampling simulations, defined as the RMSD difference of backbone atoms in the Cas8 bundle relative to both the open and closed Cas8 states (Cas8 RMSD). Wavelet analysis was performed using the code written by Heidari and co-workers^18^.

## Acknowledgments

We thank Dr. Chinmai Pindi for useful discussions and the Fraser Lab for their pre-review of our manuscript (https://prereview.org/reviews/13625755), offering critical insights that greatly helped us to improve the quality of our manuscript. This material is based upon work supported by the National Institutes of Health (Grant No. R01GM141329 to G.P.) and the National Science Foundation (Grant No. CHE-2144823 to G.P.). GP acknowledges support by the Alfred P. Sloan Foundation (Grant No. FG-2023-20431) and the Camille and Henry Dreyfus Foundation (Grant No. TC-24-063). The computational studies performed here were carried out using Expanse at the San Diego Supercomputing Center through allocation MCB160059 and Bridges2 at the Pittsburgh Supercomputer Center through allocation BIO230007 from the Advanced Cyberinfrastructure Coordination Ecosystem: Services & Support (ACCESS) program, which is supported by National Science Foundation supports grants #2138259, #2138286, #2138307, #2137603, and #2138296.

## Author Contribution

ACP performed molecular simulations, analyzed data and wrote the manuscript. SS and PRA analyzed data. GP supervised computational studies, conceived this research, and wrote the manuscript with critical input from all authors.

